# Recovering membrane interaction kinetics of single molecules from 3D tracking data

**DOI:** 10.64898/2026.04.08.717195

**Authors:** Erik Lundin, Ivan L. Volkov, Magnus Johansson

## Abstract

Interactions between cytosolic biomolecules and the bacterial inner membrane are fundamental to many cellular processes, yet directly measuring their binding kinetics in living cells remains challenging. Conventional two-dimensional single-molecule tracking analyses can be insufficient, particularly when membrane association does not markedly alter the diffusion rate. Here, we present a method to recover membrane interaction kinetics from three-dimensional single-molecule trajectories in rod-shaped bacteria. Using simulated 3D tracking data, we identify membrane-associated motion by quantifying how well short trajectory segments follow the circular curvature of the cell membrane. The resulting measure is further analyzed using a hidden Markov modeling framework, enabling robust discrimination between cytosolic and membrane-bound states and capturing the dynamics of state transitions without requiring diffusion-rate changes or direct colocalization with membrane markers. This work establishes a general framework for extracting membrane interaction kinetics from 3D single-molecule tracking data in live bacteria, and highlights the value of realistic microscopy simulations for quantitative interpretation and systematic bias assessment.

## INTRODUCTION

The cell envelope is a critical structural and functional component of the bacterial cell, with more than one third of the proteins in *E. coli* being targeted to this compartment^1^. As the cell grows, the membranes are expanded and new proteins need to be inserted in a constant flux. Mechanisms of membrane protein insertion pathways have been extensively studied over the years in reconstituted systems with purified components (see review^2^). This approach however lacks the geometrical context of the living cell. For example, in bacteria, in both co-translational (SRP pathway) and post-translational membrane protein targeting (SecA/SecB pathway), protein complexes need to be delivered to the SecYEG translocon for insertion or translocation by passive diffusion. Reconstituting these processes *in vitro* is challenging.

Single-molecule tracking microscopy opens a door to direct *in vivo* measurements of molecular interactions by following the diffusional movement of single labelled components in live cells. Though molecules typically move through the cell in 3D space, in practice, most studies rely on 2D projection (XY-plane) of the diffusion trajectory to extract transitions between different binding states. This is mainly due to difficulties in extracting the off-plane component (Z dimension) from a Gaussian point spread function (PSF) of a single emitter at high enough temporal resolution. For molecules interacting within the cytosol of bacteria, this 2D tracking approach has, however, proven very powerful to test and generate new hypotheses regarding molecular binding kinetics directly inside living cells^3–5^. For molecular interactions involving both membrane and cytosolic bindings, on the other hand, projection of diffusion trajectories onto a 2D plane leads to difficulties. That is, for rod-shaped bacterial cells oriented parallel to the focal plane, it is not possible to know whether molecular interactions occur by the membrane on a single-trajectory basis. Only cumulative analyses based on projections of the spatial distribution of these interactions are possible, hampering direct binding kinetics measurements. See for example our previous single-molecule tracking study of SRP mediated co-translational insertion of membrane proteins^3^.

It should be noted that for a single protein, membrane binding can often be registered by existing analyses. In this case, a significant change in diffusion rate upon membrane attachment can be used to resolve the kinetics based on 2D tracking data. We have previously found, however, that this becomes challenging for bigger macromolecules. For example, it turned out to be very difficult to resolve membrane binding for ribosomes involved in synthesis and targeting of inner-membrane proteins – due to only subtle changes in diffusion rate upon interaction with the membrane^3^, and the fact that translating ribosomes in general demonstrate a continuum of diffusional states^6^. Moreover, analysis of ribosomes diffusing on the curved membrane with 2D microscopy is problematic due to topological problems – some of the movements will occur perpendicular to the focal plane, thus resulting in diffusion coefficients measured from 2D projection not being representative. Hence, detection of membrane association of ribosomes in bacteria requires monitoring of its diffusion in 3D space.

A number of methods have been suggested for capturing the vertical component of a particle trajectory (see recent review^7^ for details). For example, the third dimension can be encoded in the shape of the point spread function (PSF) of the emitter by inserting an additional optical component in the emission path of the microscope, breaking the symmetry of the PSF with respect to the focal plane. The classical example of this approach is to use a cylindrical lens producing astigmatism^8^, where the PSF becomes ellipsoidal when the emitter moves out of the focal plane. Another example of PSF engineering for 3D emitter localization is the double helix PSF (DH-PSF)^9–11^. DH-PSF has two lobes that rotate in the imaging plane around the fluorophore position depending on the Z position of the emitter, making it appear as a double helix along the Z direction. It has been demonstrated that, compared to astigmatism, the Cramer-Rao bound estimated localization precision of DH-PSF is nearly constant over different Z, and on average outperforms other methods over a range of 2 µm^12^, *i*.*e*., in the length scale of bacteria. DH-PSF 3D single-molecule localization microscopy was employed in a number of studies of *E. coli* cells, including single-molecule tracking of membrane-bound proteins^13,14^, but to the best of our knowledge, the kinetics of molecular interactions has never been estimated from such 3D tracking data.

In the current study we developed a strategy for analysis of 3D single-molecule trajectories for quantification of membrane interactions kinetics in live bacterial cells, not relying on the diffusion rate measurements *per se*, but by exploiting simple geometrical properties of rod-shaped bacteria. The method was tested using simulated microscopy data obtained with a DH-PSF for molecules diffusing in an *E. coli* geometry and interacting with the membrane without an accompanying change in diffusion rate. We investigated a number of underlying kinetic models to estimate the precision and the limits of our method, and we demonstrate that this approach can be used to extract binding kinetics from 3D trajectories of membrane-interacting components in bacterial cells.

## RESULTS

### Detecting membrane trajectories by fitting a circle arc

Our suggested data analysis method for detection of membrane-associated molecules relies on the geometrical property of the bacterial membrane. In our simple model, rod-shaped bacteria such as *E. coli*, can be approximated by a cylinder with round caps. When considering only the cylindrical part of the cell, *i*.*e*., excluding the polar regions, the cell plane perpendicular to the long cell axis (YZ-plane) thus represents a circle of fixed radius. We then expect a membrane-associated molecule to follow the membrane curvature while moving, thus creating a circular path in the YZ-plane to which a circle arc should fit well. In contrast to this, a molecule diffusing in the cytosol is unlikely to follow a circular path, and its trajectory should be poorly described by a circle in the YZ-plane (Fig. 1a).

**Figure 1.**
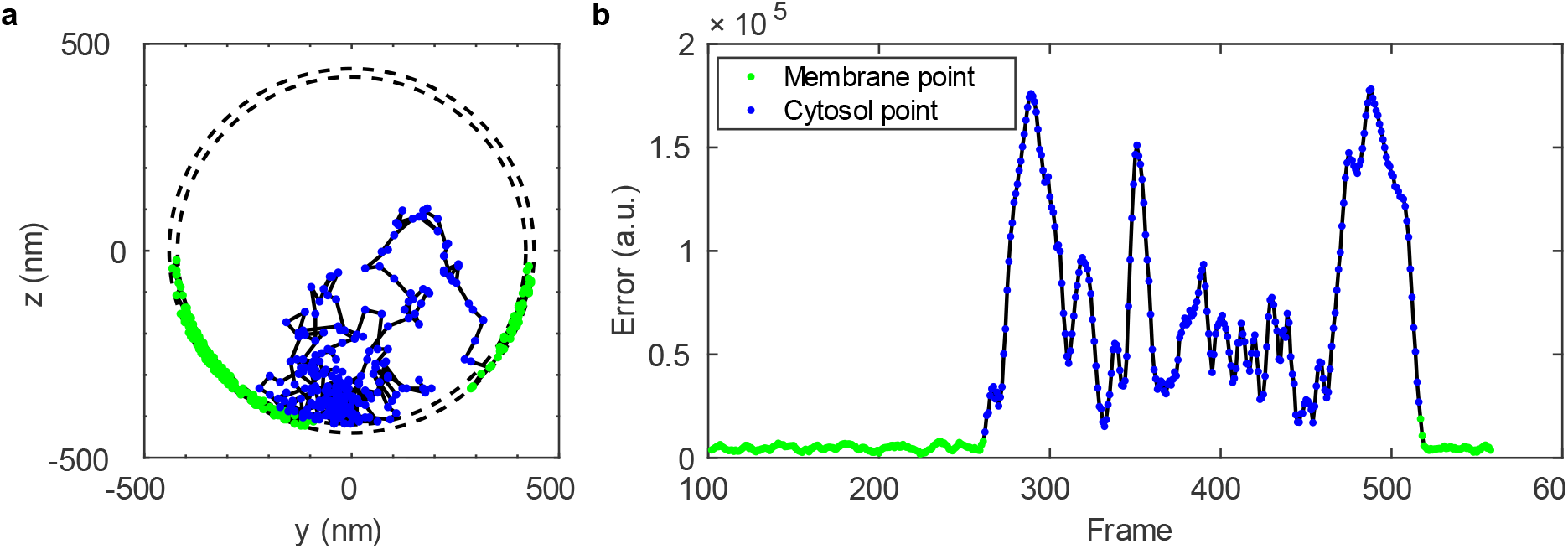
(**a**) Example trajectory of a particle diffusing in the cylindrical part of a cell (YZ projection) obtained from a reaction-diffusion simulation with trajectory points color-coded according to the states. (**b**) Error function computed with a sliding window *L* = 5 for points in the example trajectory, colored according to the states.

As the diffusion trajectories might contain transitions between membrane associated and free cytosolic movements, we defined a sliding window of *L* steps centered at each trajectory point used for calculation of a circle error function (Equation 1), where *L* is assumed to be odd. Thus, each trajectory step (except the beginning and end of the trajectory) can be associated with a computed CFE (Circle Fit Error) value (Fig. 1b).

The circle error function for a specific trajectory point, *s*, with a sliding window of *L*, is defined as

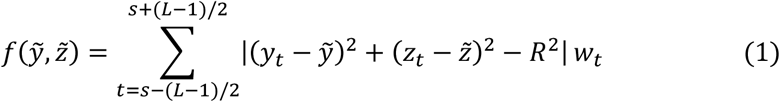

where 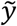 and 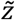 are centre position coordinates of the circle, *y*_*t*_ and *z*_*t*_ are y-and z-coordinates for step *t* in the trajectory, *R* is the radius of the cell, and *w*_*t*_ is a normalized weight function of the point. The weight function is inversely proportional to the Euclidian distance of the Y and Z localization errors of the point. Thus, points with large localization error contribute less to the total error function.

To test and validate the suggested analysis approach, we constructed simulated data where the ground-truth is known. The reaction-diffusion model in a cell geometry consists of two primary molecular states – freely diffusing in the cytosol compartment, *i*.*e*., the cytosol state (C); or diffusing in the thin membrane compartment, the membrane state (M) (Fig. 2a). To make the problem more general, the apparent diffusion rates for both states were set to be similar (0.05 µm^2^/s and 0.066 µm^2^/s), so that these states cannot be distinguished by analysis based on diffusion rate alone. The conversion rates between the states were initially set to 0.5 s^-1^, resulting in an exponentially distributed dwell time of 2 s on average for the M state, and slightly longer, 2.4 s, nearly exponentially distributed dwell time for the C state due to partially diffusion-limited conversion rate of cytosolic particles to membrane ones (Fig. S1a, b). Trajectories of 200 particles simulated for 100.1 s with a time step of 10 ms were obtained using the MesoRD^15^ software. An example trajectory from the simulation is shown in Fig. 2b.

**Figure 2.**
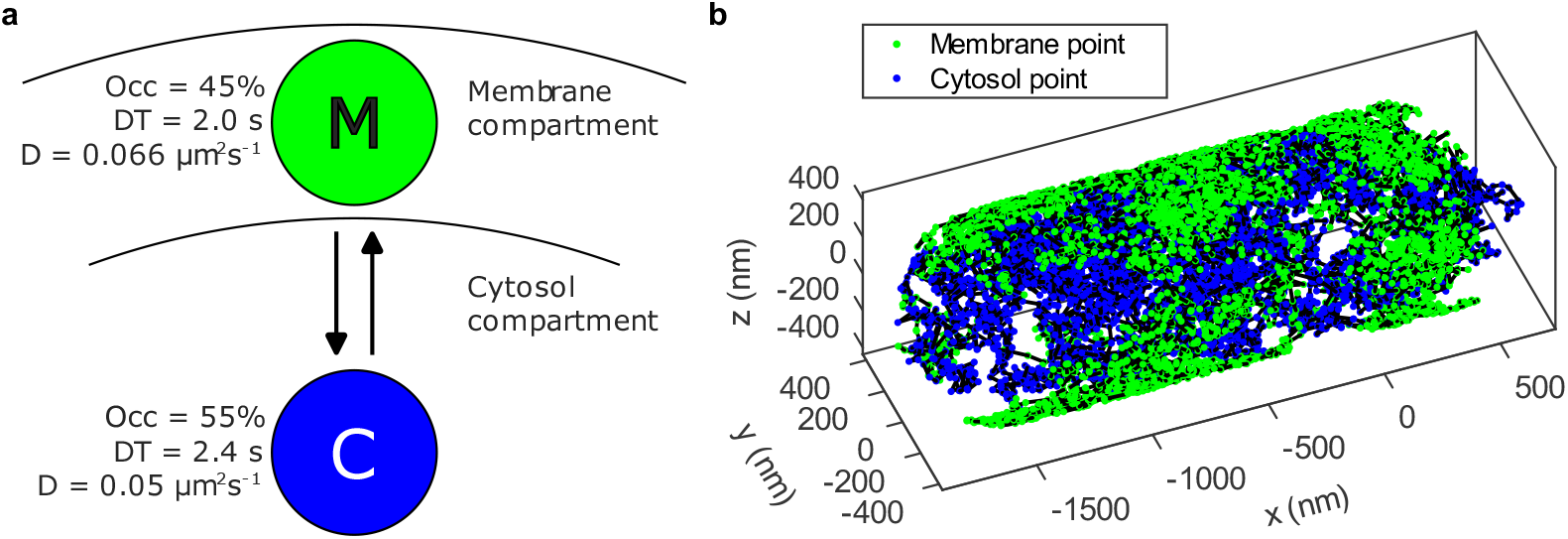
(**a**) Simulated reaction-diffusion model containing two compartments, the membrane and the cytosol, between which the particles transition. Momentary occupancy (Occ), dwell times (DT) and diffusion rate (D) for both membrane (M) and cytosolic (C) states are shown. (**b**) Example trajectory simulated from the reaction-diffusion model shown in panel a.

### Applying circle fit approach with increasing complexity of the data

#### Analysis of ground-truth trajectories

To evaluate whether the circle fit approach has any bearing, we started by analyzing simulated reaction-diffusion trajectories by computing the error function (Equation 1) with sliding window *L* = 5 at each trajectory point, with the coordinate system centered in the middle of the cell. These simulations contain no localization errors and therefore the normalized weight function, *w*, was not included. The analysis was based on only the cylindrical part of the cell by excluding segments of trajectories where the particles resided in the polar regions. The error function was computed for each trajectory point (except beginnings and ends) and the result is presented as a histogram for each ground-truth state separately in Fig. 3a. From the histogram we find that the two states can be well separated based on the value of the error function by simple thresholding. A threshold was selected to minimize the total misclassification rate, which could be as low as 0.4% in this idealized case (Fig. 3a).

**Figure 3.**
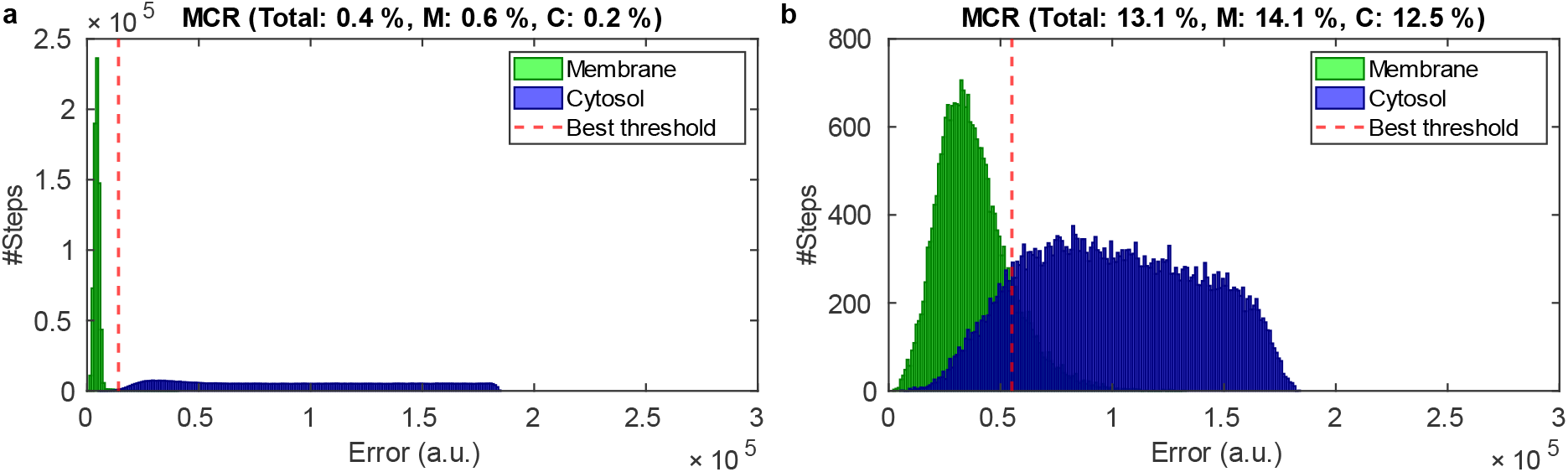
Histogram of error function values computed for individual trajectory points from the reaction-diffusion simulation (**a**) and from simulated microscopy data with realistic localization error and trajectory length (**b**). The trajectory points are classified as belonging to membrane or cytosol compartment according to the ground truth in the simulation model. The vertical dashed line represents the threshold minimizing the total misclassification rate (MCR). Total, membrane state (M), and cytosol state (C) MCR for the selected threshold are also shown.

#### Addition of localization error using simulated microscopy

As circle-error-function analysis of idealized simulated trajectories showed promising results in differentiation of membrane-bound and cytosolic diffusing particles, we proceeded to more realistic data. For this we simulated microscopy movies from existing diffusion trajectories using the SMeagol software^16^ with a realistic DH-PSF, sample background, and camera noise (Fig. S2). The frame rate was chosen to be 10 Hz, *i*.*e*., 100 ms between consecutive frames, and fluorophores were stroboscopically illuminated with 3 ms pulses to limit the diffusional blur. Fluorophore bleaching kinetics and brightness were adjusted so that tracked simulated trajectories have trajectory length and localization error in line with typical *in vivo* microscopy experiments^3,4,17^ (Fig. S3). The positions of the fluorophores in the simulated movies were then extracted with a deep-learning network model based on FD-DeepLoc^18^. In addition to X, Y, and Z positions of the fluorophore, the dot localization model also estimates the localization error, which for our data were on average 28 nm for X, 33 nm for Y, and 61 nm for Z coordinates (Fig. S3b). From the list of estimated fluorophore positions in each frame, diffusion trajectories were built using the u-track^19^ algorithm, allowing gap closure in case of missing fluorophore positions. In the resulting trajectories, we computed the circle error function as we did for the idealized reaction-diffusion trajectories above (Equation 1), but with *w*_*t*_ weights inverse proportional to the localization errors for each point. As seen from the histogram of the circle fit errors (Fig. 3b), upon introduction of realistic localization errors, both the M and the C states have significantly wider distributions of error function values, resulting in substantial overlap of the M and C components. If we again choose a threshold to minimize the misclassification, a total misclassification rate of 13.1% is theoretically achievable, compared to 0.4% for trajectories without localization errors (Fig. 3a).

#### Addition of XYZ cell origin position ambiguity using simulated microscopy

The previous simulations are still missing an important experimental error source. In a real microscopy experiment, the cells growing on, *e*.*g*., an agarose pad are never perfectly aligned with respect to the horizontal plane, generating a random offset of the cell center in the Z direction. Moreover, cell boundaries are typically detected by segmenting from phase contrast images, giving rise to imperfections in the X and Y directions (X and Y random offset). To mimic this situation, we performed the same microscopy simulations as described above, but now introducing these XY and Z offsets of 40 nm normally distributed in XY and Z on a trajectory basis.

When calculating the circle-error function on these new set of trajectories (with introduced random XY and Z offsets) based on the non-shifted, initial cell origin position, we find an even more significant overlap between cytosolic and membrane components (Fig. 4a). In this case, the best threshold provides 19.2% for the total misclassification rate (compared to 13.1% in the case without cell offsets (Fig. 3b)). The individual membrane and cytosolic trajectory points have an overall 21.9% and 17.4% misclassification rate, respectively. That is, more membrane trajectory points are misclassified as cytosolic, than *vice versa* (Fig. 4f).

**Figure 4.**
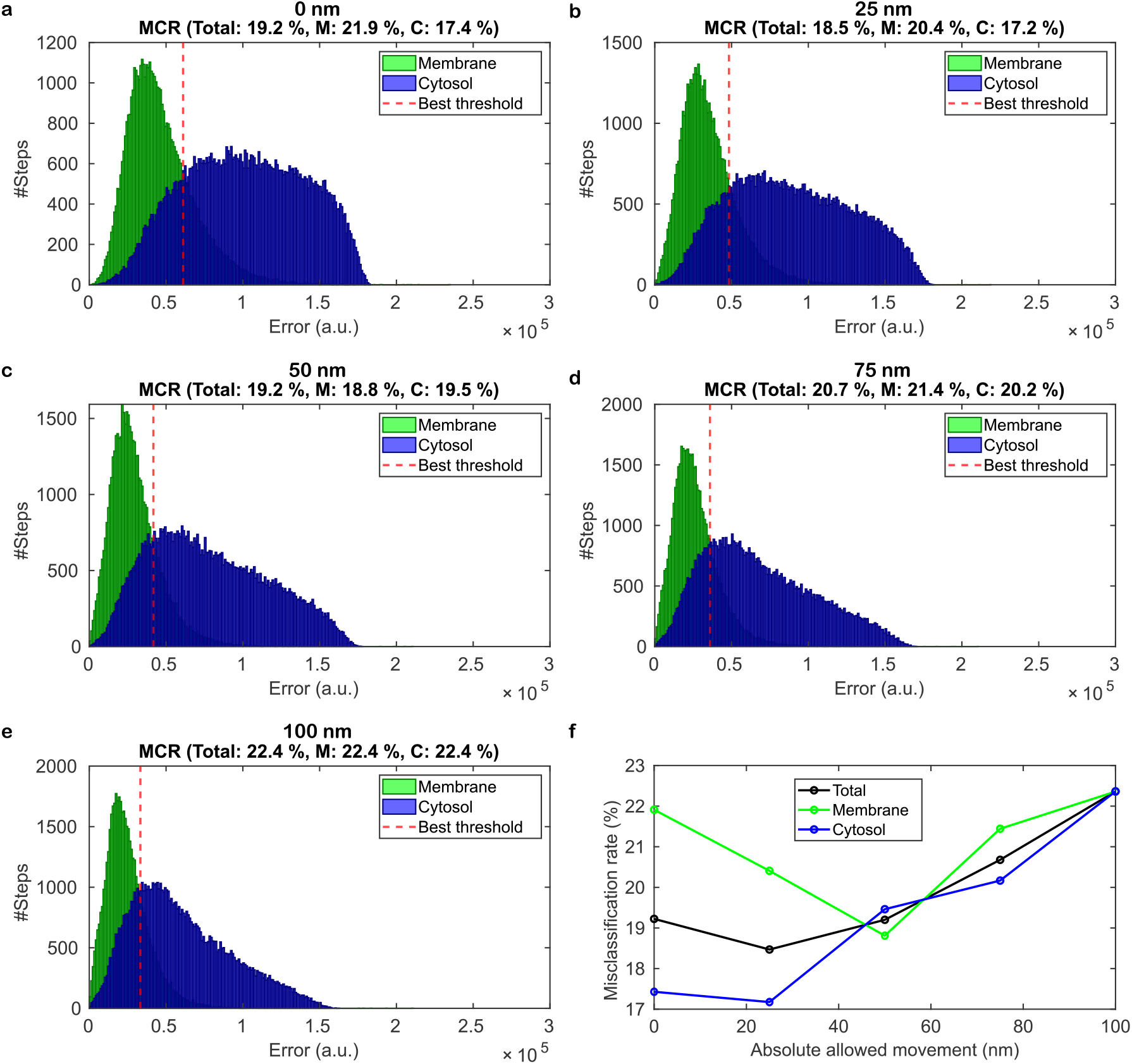
Histograms of circle-fit errors computed for individual trajectory points from simulated microscopy data with realistic localization error and trajectory length, and including normally distributed 40 nm XY and Z random offsets, mimicking imperfection of real microscopy data. The trajectory points are classified as membrane or cytosol according to the ground truth from the simulation model. Vertical dashed lines represent the threshold minimizing the total misclassification rate (MCR). Total, membrane state (M) and cytosol state (C) MCR for the selected threshold are also shown. The circle was allowed to move 0 (**a**), ±25 (**b**), ±50 (**c**), ±75 (**d**), and ±100 nm (**e**) during the fitting procedure. (**f**) Dependence of the misclassification rate on absolute allowed moving distance for the circle.

As the ground truth cell origin position in YZ plane is now shifted from (0, 0) to mimic the imperfection of *in vivo* experiments, we attempted to make our method self-correct for this by allowing a movement of the fitting circle individually for each sliding window position. In effect, we fit a circular arc to 5 trajectory points individually by minimizing the function in Equation 1, with 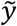 and 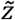 as free parameters to minimise the circle-fit error (CFE). This circle-fit procedure was then performed on the trajectories using different allowed circle-center movements of ±25, ±50, ±75 and ±100 nm in Y and Z. From the results presented in Fig. 4b-e, we find that the allowed freedom in circle fit narrowed the distribution for membrane trajectory points and changed the shape of the distribution for the cytosolic trajectory points in favor of smaller fitting errors. The total misclassification rate slightly improves, with an optimum obtained with ±25 nm allowed movement, to about 18.5% compared to 19.1% without circle movement. The most balanced individual misclassification rates with 18.8% and 19.5% for membrane and cytosolic points respectively were found with ±50 nm allowed movement, compared to 21.9% and 17.4% without circle movement (Fig. 4a, c, f). Increasing the allowed moving distance further resulted in deterioration of the total misclassification rates, up to approximately 22% for ±100 nm (Fig. 4e). We thus find that allowing movement of the circle center during the fitting procedure is a useful strategy to get better distinction between C and M states, which should however be applied with caution as too large movement makes the classification performance worse.

### Recovering of membrane binding kinetics by HMM fitting of circle fit error

So far, we have found that the circle fit approach allows us to distinguish membrane and cytosolic trajectory points with some confidence and thus can provide an estimate of the relative proportion of M and C states. However, this analysis will not provide any information on the dynamics of membrane binding. Based on the properties of the computed circle-fit error, alternating between high and low in synchrony with membrane binding events of the particle (see Fig. 1b), we reasoned that the dynamics of the process should be extractable using HMM fitting, with circle-fit error as the input (CFE-HMM). The HMM fitting procedure outputs a model of fixed size with discrete states, characterized by a CFE value, occupancy, and transition probabilities between these states. Following our previously developed approach for HMM analysis of transitions between states based on diffusion rate^3^, we analyzed the CFE trajectory data using a multi-state (8-state) HMM to account for non-exponentially distributed dwell times^6^, with subsequent coarse-graining down to two states (C and M) with a threshold chosen to minimize the misclassification rate. In this set of experiments, we used data including all sources of noise discussed above (localization errors and random XYZ offsets of trajectories) and investigated the HMM performance on CFE obtained with different circle moving distances: 0, ±25, ±50, ±75, and ±100 nm. To coarse grain 8-state models to 2 states (M and C), thresholds were selected based on a separate analysis of an additional dataset where molecules are diffusing in only M state or only C state, *i*.*e*., without state transitions (Supplementary Note 1). A lowest misclassification rate of 17.8% was obtained with ±50 nm allowed circle movement (Table 1), thus, similar to the misclassification rates obtained from simple thresholding of CFE (Fig. 4).

**Table 1.** Ground truth (GT) and CFE-HMM derived occupancies, dwell times and misclassification rates obtained for 0 - ±100 nm allowed circle movements. Errors are obtained from HMM boot-strap analysis.

|  | M occ., % | M dwell time, s | C occ., % | C dwell time, s | Misclass. rate, % |
| --- | --- | --- | --- | --- | --- |
| GT, reaction-diffusion trajectories | 45.0 | 2.0 | 55.0 | 2.4 |  |
| GT, tracked trajectories | 41.1 | 2.0 | 58.9 | 2.8 |  |
| 0 nm | $43.3 \pm 0.6$ | $8.6 \pm 0.4$ | $56.7 \pm 0.6$ | $9.9 \pm 0.4$ | 18.7 |
| $\pm 25$nm | $51.3 \pm 0.5$ | $5.7 \pm 0.2$ | $48.7 \pm 0.5$ | $6.6 \pm 0.2$ | 19.9 |
| $\pm 50$nm | $34.1 \pm 0.6$ | $5.3 \pm 0.3$ | $65.9 \pm 0.6$ | $9.9 \pm 0.4$ | 17.8 |
| $\pm 75$nm | $30.2 \pm 0.7$ | $9.2 \pm 1.1$ | $69.8 \pm 0.7$ | $12.5 \pm 0.9$ | 19.0 |
| $\pm 100$nm | $36.0 \pm 0.6$ | $6.0 \pm 0.3$ | $64.0 \pm 0.6$ | $10.9 \pm 0.5$ | 18.9 |

An important feature of the HMM fitting is that it treats the transition frequencies between the hidden states as fitting parameters. Hence, from the analysis, we directly achieve estimates of the kinetics of the system (Table 1). For ±50 nm circle movement, giving the minimal misclassification rate (17.8%), the detected dwell times are 5.3 ± 0.3 s for the M state and 9.9 ± 0.4 s for the C state, compared to 2.0 s, and 2.4 s ground truth from reaction-diffusion model for M and C dwell times, respectively. The relative occupancies in the two states are, at the same time, off by approximately 11% units. ±25 nm circle movement gives better approximation of C dwell time (6.6 s) and occupancy (off by 6% units). Thus, the choice of allowed circle-moving distance can be dictated by the importance of the parameter to be extracted. Ground-truth occupancies and dwell times computed for annotated tracked trajectories are only marginally different from ground-truth parameters of the reaction-diffusion trajectories (Table 1). Hence, tracking artefacts do not contribute significantly to the observed mismatch between CFE-HMM and ground truth from the underlying reaction-diffusion model. An example trajectory demonstrating membrane binding along with calculated CFE and CFE-HMM fits for ±25 nm circle movements is shown in Fig. 5.

**Figure 5.**
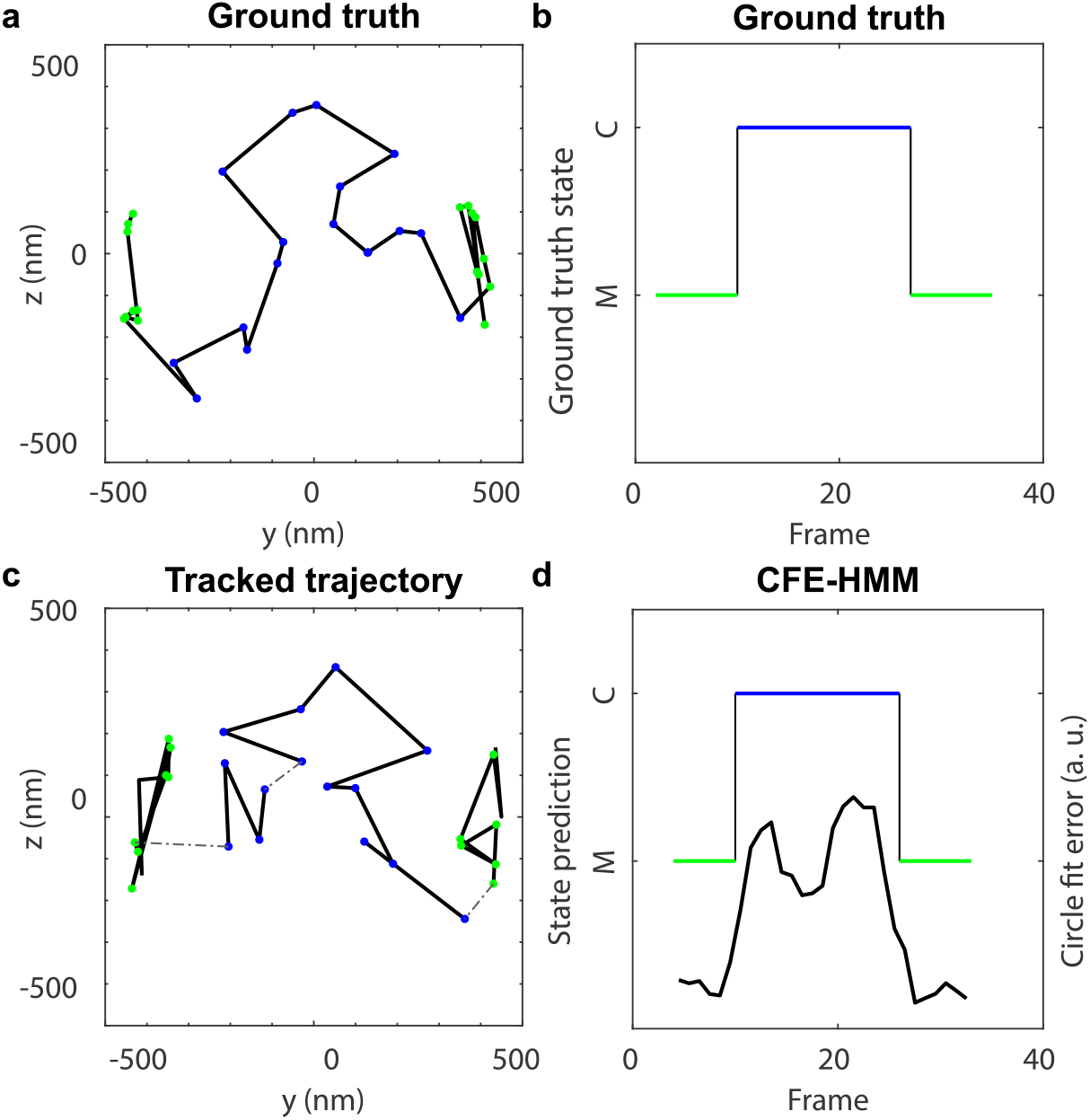
Ground truth trajectory (**a**) and corresponding trajectory obtained by tracking simulated microscopy with localization errors, missed trajectory segments (dashed lines), and cell origin XYZ offsets (**c**). Trajectory points are color-coded according to the selected state. Ground truth states (**b**) and corresponding CFE curve along with CFE-HMM predicted states obtained with ±25 nm allowed circle movements (**d**).

### Exploring membrane binding in different kinetic models by CFE-HMM

The underlying kinetic model for the data discussed above is based on approximately 2 s nearly exponentially distributed dwell times for both M and C states. As circle fitting operates on a sliding window of 5 trajectory points, and data is sampled with 100 ms time intervals, we suspect that some of the short events might be “smoothed” by the sliding window (*i*.*e*., 500 ms) and, thus, not detected. This would lead to overestimation of dwell times – something apparent in all our fittings (Table 1). Indeed, by aligning stretches of the same ground-truth state (1 – 20 frames long) to the CFE-HMM prediction, we found that the percentage of well-predicted trajectory stretches drops sharply when the stretch length is shorter than 5 (Fig. S5). At the same time, from our previous analyses of single-molecule tracking data^6,17^, we also know that very long dwell times are more difficult to measure precisely. That is, due to rareness of transition events, there is a relatively higher contribution of analysis and tracking artefacts, which gives rise to false transition events. Hence, to explore a wider range of kinetics, we applied CFE-HMM analysis to simulated tracking data generated with exponentially distributed dwell times from 1 s to 8 s in each state. Further, as many biological processes are multi-step reactions and, thus, their dwell times are not exponentially distributed, we also explored models where only the M state or both the M and the C states have gamma distributed dwell times from 1 s to 8 s (Fig. S1, Supplementary Note 2). The analysis was performed in a similar manner as above. Here we discuss the results obtained with ±25 nm circle movement as a consensus moving distance providing the dwell times closest to the ground truth (Table S1). We find that CFE-HMM is capable of detecting the difference between models with 1, 2, 4, and 8 s dwell times, but systematically overestimate the membrane-bound dwell time by 2-3 s on average (Fig. 6b). Models where one or both of the states have gamma-distributed dwell times are on average recovered better than models with exponential dwell times. As discussed above, this is most probably due to missed short events which are more prevalent for exponentially distributed times (Fig. S1). Occupancies obtained for different models differ from ground truth by about 3-7 % units (Fig. 6a). Dwell times for the C state for 1-4 s models suffer from similar overestimation as for M state (Fig. 6c). For models with 8 s dwell times, HMM fitting did not converge with respect to the C-state dwell time (discussed below), thus making cytosolic dwell times longer than approximately 4 s undetermined. Because our primary objective is to characterize the kinetics of the membrane-bound state, however, we do not consider this to be a significant concern.

**Figure 6.**
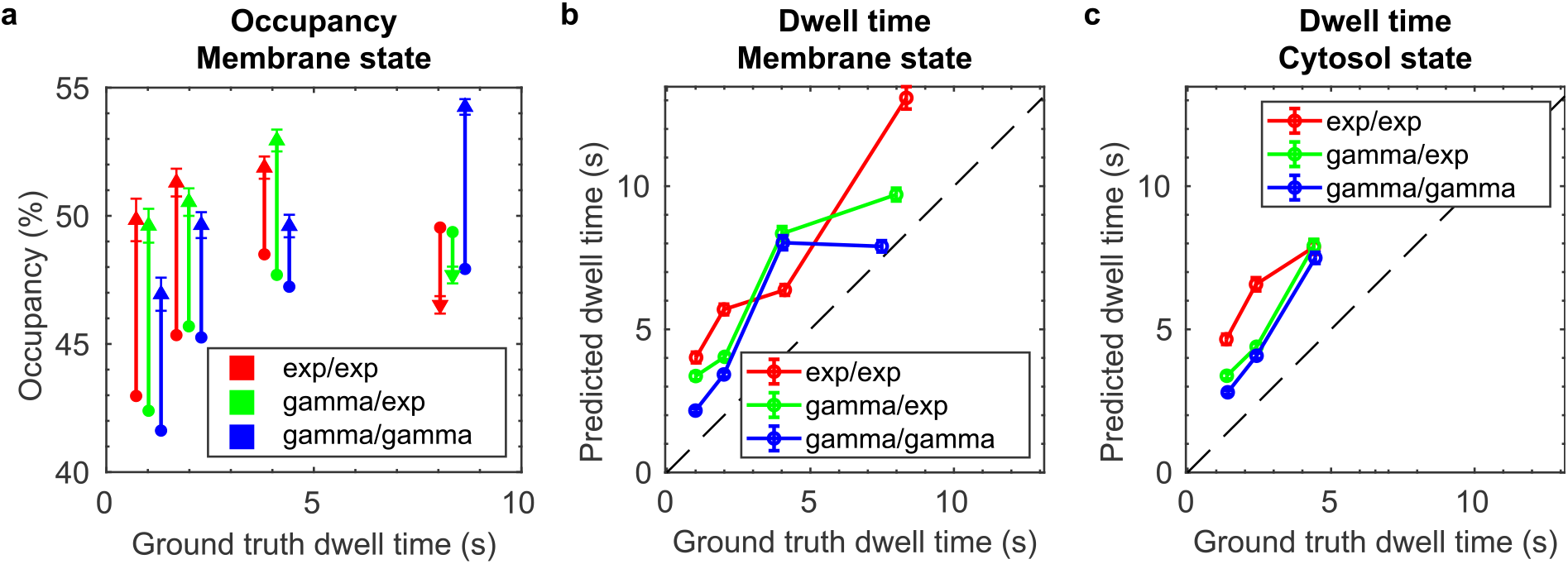
Occupancy prediction (triangles) relative to ground truth occupancy (circles) (**a**) and predicted dwell time (**b, c**) plotted against the ground truth dwell time of membrane state (**a, b**) and cytosol state (**c**) for models with dwell time of 1, 2, 4, and 8 s for membrane and cytosol state, with 8 s model only for membrane state (**a, b**). The membrane and cytosol state dwell time were either both exponentially or gamma distributed, or the dwell time of the membrane state gamma distributed and the cytosol state exponentially distributed. Error bars represent boot-strap errors.

### Exploring the robustness of the method

From the analysis above we find that dwell times are typically best recovered from models with gamma-distributed dwell times in both C and M states. Those models are probably also best describing *in vivo* biological processes related to molecular interactions with the membrane (*e*.*g*., co- and post-translational protein insertion), as such processes are likely multi-step reactions. We finally wanted to investigate the robustness of the CFE-HMM method with respect to available amount of data as well as parameters used for analysis.

A machine-learning algorithm, such as HMM, is sensitive to the amount of data available for analysis. With regards to this we investigated convergence of the HMM fitting for models with gamma-distributed dwell times (Supplementary Note 3). We found that models with shorter dwell times (more frequent transitions) converge faster than models with longer dwell times (more rare transitions) and that state occupancy typically converges quicker than the dwell time parameter (Fig. S7). Typical relative standard deviation (RSD) of the converged occupancy is below 3% for 1-4 s models and below 12% for 8 s model (Table S2). The RSD of dwell time is below 11% and similar to HMM-derived relative boot-strap error (Table S1) for all the models, except C state in 8 s model, which converged much more poorly, only down to 56% RSD (±12 s STD). From these results we can conclude that all fitted parameters in this study have converged, except for the C-state dwell time in the 8 s model, which showed poor convergence even with 400 000 trajectory steps (Fig. S7). Those numbers were, hence, not reported in the figures. The M-state dwell time for the 8 s model did, however, converge after approximately 10 000 trajectory steps.

Further, we investigated how sensitive the detected dwell times are to the threshold selection for coarse-graining multi-state HMM to M and C states in the models with gamma distributed dwell times. Variation of the threshold by ±10 and ±20% resulted in RSD for detected dwell times not exceeding 20% (Fig. S8a, b and Table S3).

Finally, we also tested how the estimated dwell times depend on the selected circle movement by allowing ±50 nm or no movement in addition to the optimal ±25 nm movement (Fig. S8c, d). In this case, the M/C classification threshold was optimised for each moving distance. We found that dwell times have RSD up to 48% in some models, thus making the allowed circle movement the most sensitive parameter in our analysis. This parameter should, hence, be chosen with caution.

## DISCUSSION

In the current study we present a method for detection and quantification of interactions of cytosolic biomolecules with the membrane in rod-shaped bacteria, and demonstrate its capability to derive dwell times and occupancies of membrane-associated and cytosolic states. First, we established the general principle for the membrane-binding analysis in case of particles diffusing in the cytosol and interacting with the cell membrane without an observable change of the diffusion coefficient. To mimic *in vivo* microscopy data, we gradually increased the complexity of the model from long ground-truth trajectories to photobleaching-limited trajectories with realistic localization errors in cells with random offsets in X, Y and Z.

To determine whether the molecule is associated with the curved bacterial membrane or freely diffusing in the cytosol, we exploited a simple geometrical property of the rod-shaped bacteria – membrane-bound molecules are likely to follow the membrane curvature while diffusing. To detect this diffusional property, we performed a circle fit on trajectory points within a sliding window, and calculated the corresponding circle fit error (CFE) with the presumption that membrane trajectory points likely result in lower CFE, compared to cytosolic trajectory points. Thus, the method does not exploit direct colocalization of the molecule of interest with the membrane. In addition, this analysis does not rely on a change in the diffusion coefficient, but only depends on how well a particle trajectory follows the circular arc of the cell membrane in the YZ plane. We found that for long ideal trajectories without localization error, almost perfect classification of membrane and cytosolic trajectory points can be reached (0.4% total misclassification rate), while in a more realistic case, with trajectories containing localization error and XYZ offset, the misclassification rate is higher (about 19%).

To extract binding kinetics from the diffusion trajectories, CFE values assigned to every trajectory point were fitted by multi-state CFE-HMM, where the states were subsequently classified as membrane-bound or cytosolic based on separate reference simulations. We found that the CFE-HMM analysis is capable of detecting dwell times of membrane-associated and cytosolic states. Best performance with respect to dwell-time precision was achieved for models with gamma-distributed dwell times, which we also perceive as most biologically relevant. Typically, the method overestimate dwell times by 2-3 seconds on average. However, we expect that in most cases, the exact bias can be inferred by analyzing simulated microscopy data of the reaction-diffusion model under study^3^.

The presented method is currently limited to only the cylindrical part of the cell. However, a similar approach with some modifications could potentially be used to analyze trajectories within the whole cell. The presented circle-fit method further requires an initial guess of the cell origin position and cell radius. In the current simulations, we have assumed that the cell origin is known with ±40 nm precision in Y and Z, and that the cell radius is fixed. Both the cell radius and cell origin position should, hence, be provided for analysis of real microscopy data. In practice, this can be achieved by imaging the cell contour in a second imaging channel.

Finally, in the present simulations, the acquisition frame rate was selected such that trajectories of membrane-diffusing molecules get well distributed on the curved membrane during binding events, and the average step length exceeds the localization error. In a live-cell experiment, the frame rate should, thus, be tuned depending on the diffusion rates of the molecule of interest. At the same time, the frame rate needs to be fast enough to capture short-lasting events and fast transitions, in practice, introducing a trade-off in the analysis and, hence, a limitation to the approach. Further, the optimal allowed circle moving distance and threshold position for classification of M and C state in the large HMM model are additional parameters that likely depends on the data acquisition rate and localization precision.

## CONCLUSIONS

This proof-of-principle study demonstrates the strong potential of 3D single-molecule microscopy for investigating membrane-binding molecules in rod-shaped bacteria. By leveraging the inherent geometric signature of membrane curvature, we can identify membrane-binding events and infer their underlying kinetics without requiring direct colocalization of the tracked molecule with a membrane marker. We anticipate that further development of the analytical framework, particularly through the integration of AI-based pattern-recognition algorithms^20^, will enhance the method even further.

## MATERIALS AND METHODS

### Simulation of diffusion trajectories and microscopy data

Simulation of the diffusion trajectories was performed similarly to previous studies^3,17^. In short, trajectories for the defined reaction-diffusion model were simulated by MesoRD software^15^. These trajectories were used to construct simulated microscopy movies with SMeagol software^16^ and then analysed with the same image analysis pipeline as experimental microscopy data.

#### Simulating reaction-diffusion kinetics using MesoRD

The cell geometry consisted of a cytosol compartment constructed from a cylinder of length 3 µm and a radius of 0.42 µm with half-spheres of radius 0.42 µm as poles. The membrane compartment was constructed by starting with a structure like the cytosol compartment, but with the cylinder and half-sphere cylinder being 0.44 µm, then subtracting the cytosol compartment, leaving a 0.02µm layer around the cytosol compartment. A similar geometry has been used by us previously for simulations of Signal Recognition Particle (SRP) interacting with ribosomes and the membrane^3^. The thickness of the membrane compartment was, however, now reduced to achieve more realistic two-dimensional diffusion by the particles on the membrane surface.

The reaction diffusion model (Fig. S6) has two primary states, the membrane state B with diffusion coefficient 0.066 µm^2^/s and the cytosol state D with diffusion coefficient 0.05 µm^2^/s, with additional states to give the correct behaviour as explained in Supplementary Note 2.

Simulations were performed in batches, with the number of batches corresponding to the models dwell time. For example, for models with 2 s dwell times, two batches were used. One batch consisted of 12 simulations with 200 particles initiated in the cytosolic state A. First, the system was equilibrated for 5 s (simulation time) and then simulated for an additional 100.1 s to acquire reaction and trajectory data. The time step used was 0.01 s and the spatial discretization was 0.005 µm, giving a membrane layer of four cubes.

In case of added XY offsets mimicking cell segmentation imperfections, trajectories obtained from MesoRD simulation were shifted in X and Y directions with random normally distributed offsets with standard deviation 40 nm and with mean 0 nm.

#### Simulated video microscopy using SMeagol

Simulated microscopy movies were constructed with the SMeagol software^16^ using experimentally derived background and parameters giving rise to movies typical to *in vivo* microscopy. Fluorophore spots (one per cell) were simulated using an experimentally derived point spread function (PSF) constructed from a 25 nm Z-stack of a fluorescent bead, similarly to a previous PSF model^17^. The simulated microscopy movies had a frame time of 100 ms and illumination time of 3 ms. Cell segmentation masks were generated, filling internal holes with convex hull and then shrunken 240 nm with erosion^3^.

To mimic the cell-to-cell depth distribution on a microscopy sample, normally distributed 40 nm (mean 0 nm) offset in Z position of cells was added when applicable.

#### Aligning SMeagol trajectories with tracked trajectories

Trajectory-step identity (based on the reaction-diffusion model from MesoRD) is retained in SMeagol output data. This information is however lost during single-molecule tracking image analysis. To align tracked data with the ground-truth, the following procedure was performed. The Euclidian distance between a tracked trajectory point and the ground-truth coordinates was calculated, and the ground-truth trajectory giving the minimal distance was pre-selected for a given frame. The same procedure was repeated for each trajectory frame, and the ground-truth trajectory corresponding to the tracked one was selected based on majority vote from individual frames. This links ground truth and tracked trajectories, allowing state identity for each tracked trajectory to be restored.

### Data analysis

#### Dot detection and trajectory building

Image analysis was performed using a previously described MATLAB-based pipeline^17^. 3D positions of the fluorophores were detected using a pre-trained deep-learning network model based on FD-DeepLoc^21^. Dots passing a 60% confidence threshold were kept for downstream analysis.

Trajectory building was performed using the u-track algorithm^19^, connecting points in consecutive frames with a search radius of 800 nm in the XY plane (i. e., Z component was ignored). The gap filling feature allowed missed fluorophore position for one frame between well-detected trajectory segments.

#### Construction and training of the dot detection model

The dot detection model was constructed as described earlier^21^. First a spline model of the double-helix PSF was constructed from a 50 nm Z-stack of fluorescent beads using SMAP^22^. The neural network model was then trained on simulated microscopy images generated using the PSF spline model and the EMCCD camera model.

#### Circle fit

Circle-error function values were calculated according to Equation 1. The weight function, when used, was calculated by first calculating the inverse of the Euclidian distance for the Y, Z localization error for each trajectory point. This was then normalized over all points to get the weight function.

When movement in Y and Z was allowed, the circle fit was performed by minimizing the circle error function using the MATLAB function “fmincon” that minimize a multivariable function under given constraints. As a constrain, the circle origin was allowed to move ± defined distances from the initial guess independently in both Y and Z direction.

The circle radius used when calculating circle error function was set to 430 nm for all cells.

#### CFE-HMM analysis

Circle-fit errors computed for each trajectory step (except beginnings and ends) were analyzed by Hidden Markov Modeling (uncertainSPT^23^), similar to the procedure described in reference ^3^ for diffusion HMM. Doing so, we used the same HMM program as in reference ^3^ and treated CFE similarly to step lengths in the diffusion HMM. The data were first fitted to 8 states, and then coarse-grained to two states using a threshold to separate biologically relevant states (M and C). Coarse-grained occupancies were calculated as sums of the state occupancies included in the coarse-grained state, and coarse-grained mean dwell times were estimated from the coarse-grained transition matrix (Note S1 in reference ^24^).

## DATA AVAILABILITY

A detailed list of parameters for data analysis will be available together with raw data in the SciLifeLab data repository.

## AUTHOR CONTRIBUTIONS

E.L., I.L.V., and M.J. designed research. E.L. performed simulations and wrote analysis code. E.L. and I.L.V. analyzed data. I.L.V. and M.J. conceived the project. E.L., I.L.V., and M.J. wrote the manuscript.

## DECLARATION OF GENERATIVE AI AND AI-ASSISTED TECHNOLOGIES IN THE WRITING PROCESS

During the preparation of this work the authors used Microsoft Copilot in order to improve the readability of certain parts of text. After using this tool, the authors reviewed and edited the content as needed and take full responsibility for the content of the publication.

## ACKNOWLEDGEMENTS

Thea authors would like to thank Yoav Shechtman’s lab for providing a DH phase mask required to generate experimental DH PSF used for simulation microscopy. This study was made possible by grants from the ERC (947747-SMACK) and the Swedish Research Council (2023-03383, 2024-06104). The computations and data management were enabled by resources provided by the Swedish National Infrastructure for Computing at UPPMAX, partially funded by the Swedish Research Council through grant agreement no. 2018-05973.

