## Supplementary Information for "Recovering membrane interaction kinetics of single molecules from 3D tracking data"

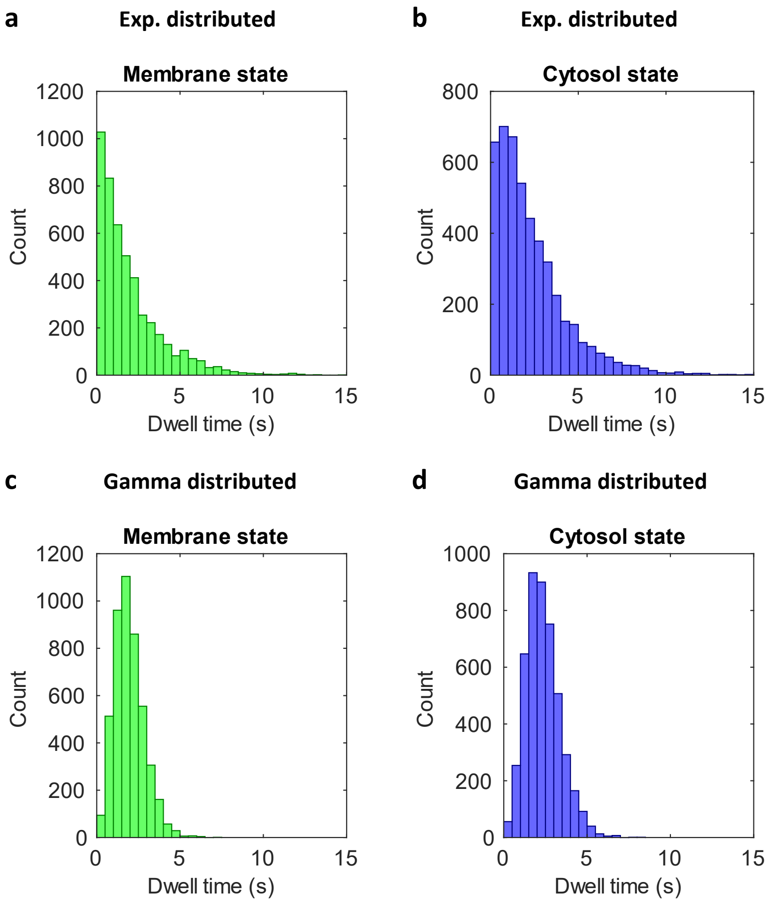

**Figure S1.** The distributions of dwell times of membrane (**a, c**) and cytosol state (**b, d**) from reaction-diffusion models with exponentially (**a, b**) and gamma (**c, d**) distributed dwell times, each approximately 2 s on average.

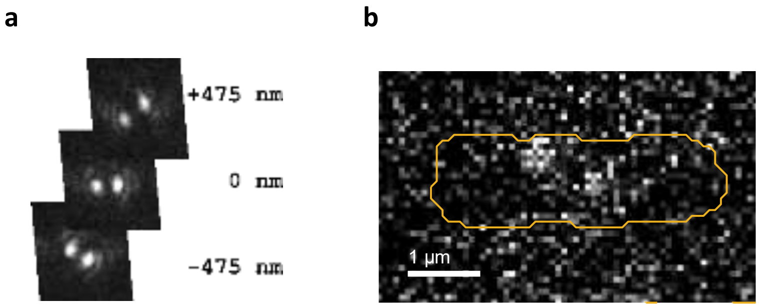

**Figure S2.** DH-PSF used for microscopy simulation, shown at three different depths (**a**), and an example of simulated microscopy movie frame with DH-PSF representing molecules labelled with fluorophores moving in an *E. coli* cell geometry (**b**).

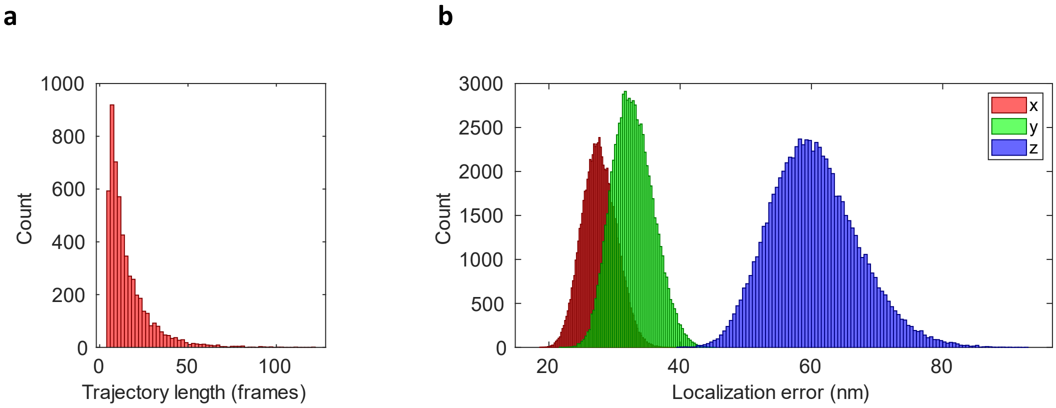

**Figure S3.** The distributions of trajectory lengths (**a**) and localization errors for x, y and z coordinates (**b**) obtained by dot detection and trajectory building algorithms from the analysis of simulated microscopy data.

### Supplementary Note 1. Selection of threshold for best separation of cytosolic and membrane states in coarse-grained HMM models

To find a threshold for a CFE-HMM 8-states model that can separate membrane states from cytosolic state, two models were simulated with only a membrane or only a cytosol state correspondingly, and diffusion coefficients corresponding to M and C states in previous models (Fig. 2, main text). The tracked trajectories from the simulated microscopy movies of the two models were analysed by CFE-HMM as one combined dataset using the same circle fit moving distances as were used for the analysis of models with transitions (0, ±25, ±50, ±75 and ±100 nm).

From the 8-state model, a list of seven potential thresholds placed between each pair of the neighbouring states was determined. For each potential threshold, the membrane, cytosol and total misclassification rate was calculated, and the threshold resulting in the minimal total misclassification rate for a given allowed moving distance was selected (Fig. S4).

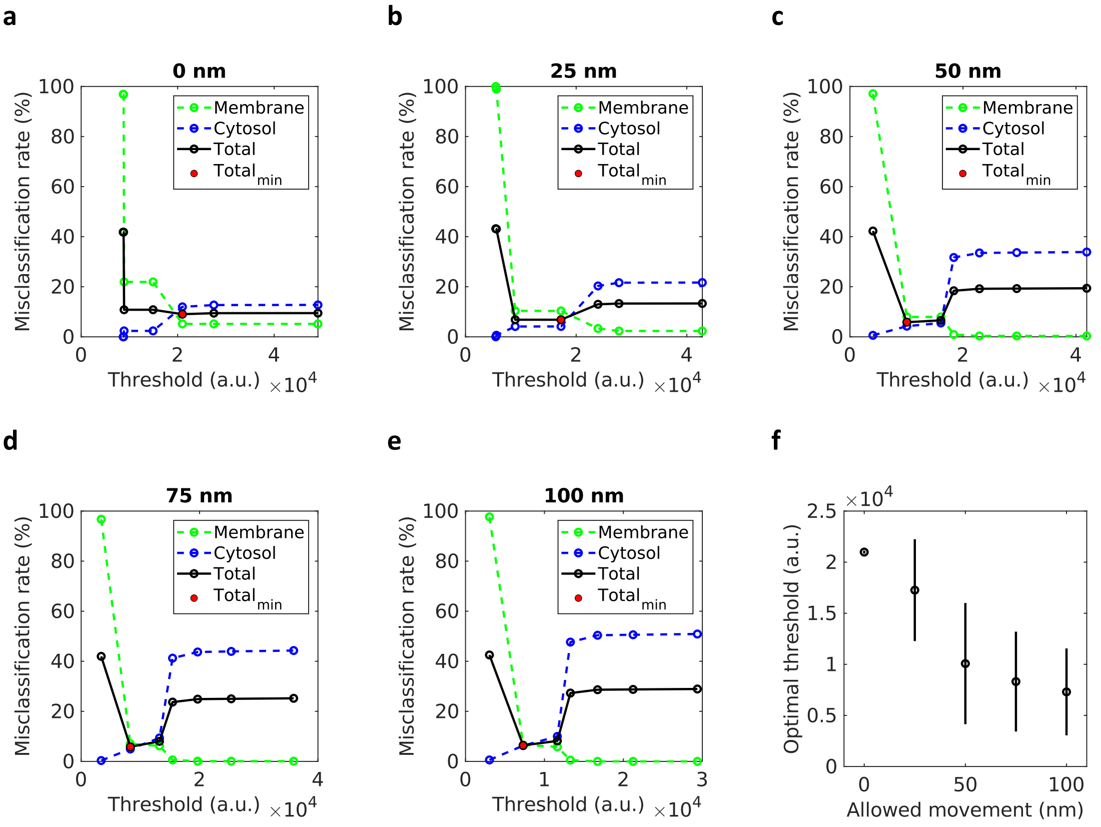

**Figure S4.** The membrane, cytosol and total misclassification rates calculated using potential thresholds for CFE-HMM with 0 (**a**), ±25 nm (**b**), ±50 nm (**c**), ±75 nm (**d**) and ±100 nm (**e**) allowed circle movement distance. The threshold value resulting in minimal total misclassification rate (red circle) is chosen as optimal threshold. Dependence of the optimal threshold value on the moving distance (**f**). For each moving distance, the selected threshold is the middle value within a range of thresholds that would result in the same coarse-grained model.

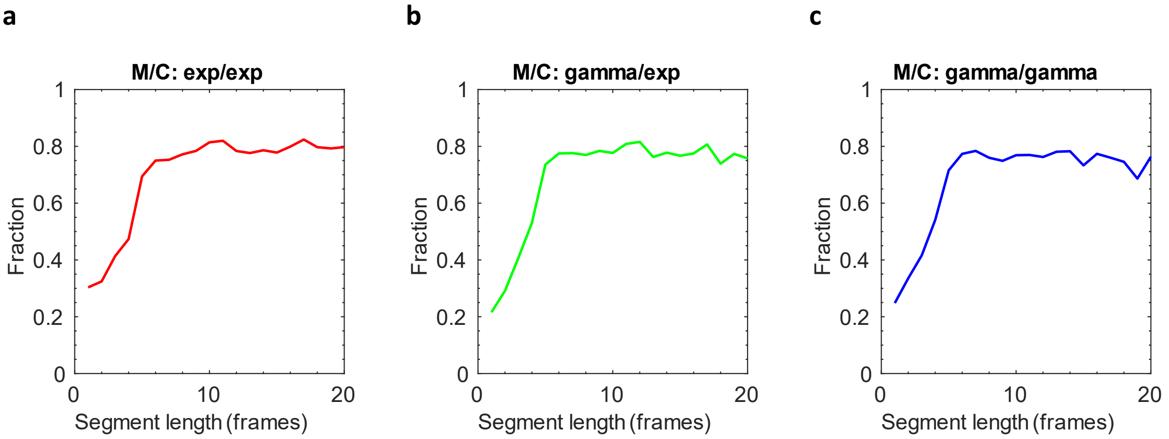

**Figure S5.** Performance of state prediction depending on trajectory segment length. Tracked trajectories were divided into segments of the same ground truth state. For each segment length (1-20 frames) the fraction of segments with more than 80% of points correctly predicted within this segment is calculated. The fraction is plotted against the segment length for models with 2 s mean dwell time, where the dwell time distribution is exponential for both the membrane and cytosol state (**a**), gamma for the membrane state and exponential for cytosol state (**b**), or gamma for both the membrane and cytosol state (**c**).

### Supplementary Note 2. Simulation of models with different kinetics

The model shown in Fig. 2 (main text) contains two primary states (M and C), with additional non-observable substates added for computational reasons (Fig. S6a). The primary M state in the simple model (Fig. 2) will have an exponentially distributed dwell time (Fig. S1a), whereas the primary C state in the simple model will have a nearly exponentially distributed dwell time (Fig. S1b), due to the delay caused by diffusion to the membrane before it can transform to the M state.

To simulate non-exponentially distributed dwell times we constructed models with sequential substates with the same diffusion coefficients and reaction rates (Fig. S6b, c). The models with sequential states have gamma distributed dwell times of the primary states (Fig. S1c, d).

First, the M state was given a gamma distributed dwell time by replacing the main M state with five sequential states with transition rates between them 5x higher than the single-step rate (Fig. S6b). This results in an M-state dwell time which is non-exponentially distributed, while the C state dwell time is still nearly exponentially distributed. Second, we also modified the C primary state to have a gamma distributed dwell time by, similar to the M state, replacing the main C state with a sequence of five equal substates at 5x higher transition rates (Fig. S6c), resulting in gamma distributed dwell times for both M and C primary states (Fig. S1c, d).

To modulate the mean dwell time from initial 2 s to 1, 4 or 8 s, rate constants (‘k’ in Fig. S6) were modified correspondingly.

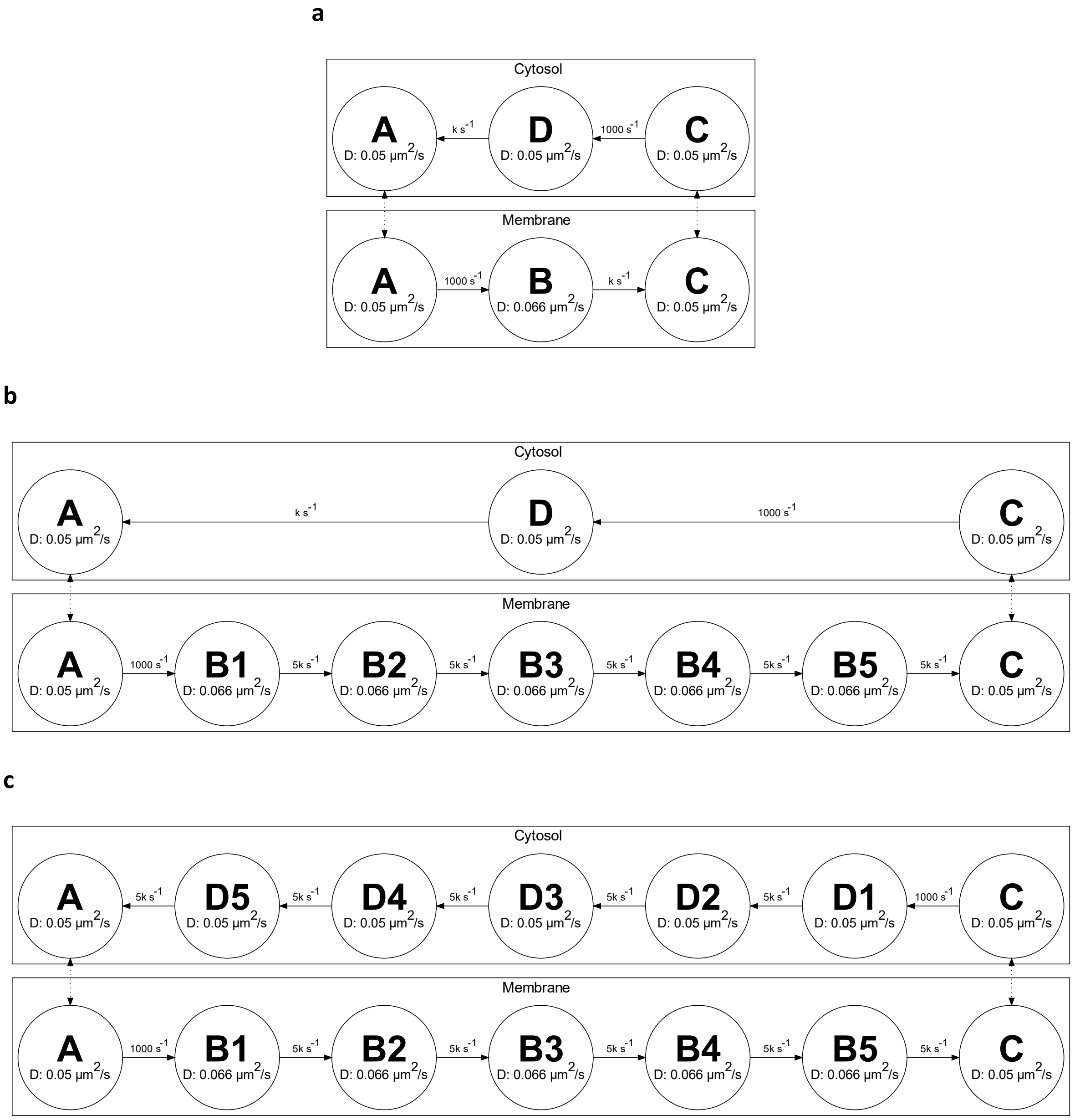

**Figure S6.** Reaction-diffusion models of membrane-cytosol dynamics simulated in MesoRD with nearly exponentially distributed dwell times for both M and C states (**a**), gamma distributed dwell time for M and exponential for C state (**b**), and gamma distributed dwell times for both M and C states (**c**). Transition rates between substates equals either k $s^{-1}$ (for exponential dwell time) or 5k $s^{-1}$ (for gamma distributed dwell time), where k = 1 for 1 s, k = 0.5 for 2 s, k = 0.25 for 4 s, and k = 0.125 for 8 s dwell time.

**Table S1.** Occupancy and dwell time, with boot-strap error and relative boot-strap error, for models with M and C dwell times of 1 – 8 s distributed as either both exponential, gamma and exponential, or both gamma. Parameters were calculated from reaction-diffusion simulations (MesoRD GT), from ground truth of tracked trajectories (tracked GT), and from CFE HMM with allowed circle movement of 0, ±25, ±50, ±75 or ±100 nm.

| M/C: exp/exp dwell time 1 s | | | | | | | | | | |
|---|---|---|---|---|---|---|---|---|---|---|
| | M occ., % | M occ. boot-strap error | M dwell time, s | M dwell time boot-strap error | M dwell time relative boot-strap error, % | C occ., % | C occ. boot-strap error | C dwell time, s | C dwell time boot-strap error | C dwell time relative boot-strap error, % |
| **MesoRD GT** | 43,0 | | 1,0 | | | 57,0 | | 1,4 | | |
| **tracked GT** | 37,6 | | 1,0 | | | 62,4 | | 1,7 | | |
| **0 nm** | 41,3 | 0,9 | 6,6 | 0,4 | 5,4 | 58,7 | 0,9 | 11,5 | 0,7 | 6,1 |
| **25 nm** | 49,8 | 0,8 | 4,0 | 0,2 | 4,6 | 50,2 | 0,8 | 4,7 | 0,2 | 3,9 |
| **50 nm** | 54,0 | 0,8 | 3,6 | 0,1 | 4,1 | 46,0 | 0,8 | 3,8 | 0,1 | 3,5 |
| **75 nm** | 57,9 | 0,8 | 3,6 | 0,1 | 4,2 | 42,1 | 0,8 | 3,7 | 0,1 | 3,8 |
| **100 nm** | 55,0 | 2,8 | 4,2 | 0,2 | 5,8 | 45,0 | 2,8 | 4,3 | 0,3 | 7,8 |

| M/C: gamma/exp dwell time 1 s | | | | | | | | | | |
|---|---|---|---|---|---|---|---|---|---|---|
| | M occ., % | M occ. boot-strap error | M dwell time, s | M dwell time boot-strap error | M dwell time relative boot-strap error, % | C occ., % | C occ. boot-strap error | C dwell time, s | C dwell time boot-strap error | C dwell time relative boot-strap error, % |
| **MesoRD GT** | 42,4 | | 1,0 | | | 57,6 | | 1,4 | | |
| **tracked GT** | 38,2 | | 1,0 | | | 61,8 | | 1,6 | | |
| **0 nm** | 46,0 | 0,8 | 9,8 | 0,8 | 8,1 | 54,0 | 0,8 | 9,5 | 0,7 | 6,9 |
| **25 nm** | 49,6 | 0,7 | 3,4 | 0,1 | 3,9 | 50,4 | 0,7 | 3,4 | 0,1 | 3,3 |
| **50 nm** | 46,3 | 1,0 | 2,6 | 0,2 | 6,4 | 53,7 | 1,0 | 1,0 | 0,0 | 2,3 |
| **75 nm** | 46,8 | 3,0 | 3,2 | 0,2 | 7,4 | 53,2 | 3,0 | 1,1 | 0,1 | 10,7 |
| **100 nm** | 55,4 | 0,8 | 3,7 | 0,2 | 4,8 | 44,6 | 0,8 | 0,8 | 0,0 | 2,2 |

| M/C: gamma/gamma dwell time 1 s | | | | | | | | | | |
|---|---|---|---|---|---|---|---|---|---|---|
| | M occ., % | M occ. boot-strap error | M dwell time, s | M dwell time boot-strap error | M dwell time relative boot-strap error, % | C occ., % | C occ. boot-strap error | C dwell time, s | C dwell time boot-strap error | C dwell time relative boot-strap error, % |
| **MesoRD GT** | 41,6 | | 1,0 | | | 58,4 | | 1,4 | | |
| **tracked GT** | 37,6 | | 1,0 | | | 62,4 | | 1,6 | | |
| **0 nm** | 41,2 | 0,7 | 2,8 | 0,1 | 4,5 | 58,8 | 0,7 | 1,4 | 0,0 | 2,4 |
| **25 nm** | 46,9 | 0,6 | 2,2 | 0,1 | 3,3 | 53,1 | 0,6 | 2,8 | 0,1 | 2,6 |
| **50 nm** | 43,5 | 0,7 | 1,7 | 0,1 | 3,5 | 56,5 | 0,7 | 1,0 | 0,0 | 1,7 |
| **75 nm** | 48,9 | 0,6 | 2,2 | 0,1 | 3,6 | 51,1 | 0,6 | 0,9 | 0,0 | 1,8 |
| **100 nm** | 60,1 | 0,6 | 3,4 | 0,1 | 3,2 | 39,9 | 0,6 | 2,5 | 0,1 | 2,9 |

| M/C: exp/exp dwell time 2 s | | | | | | | | | | |
|---|---|---|---|---|---|---|---|---|---|---|
| | M occ., % | M occ. boot-strap error | M dwell time, s | M dwell time boot-strap error | M dwell time relative boot-strap error, % | C occ., % | C occ. boot-strap error | C dwell time, s | C dwell time boot-strap error | C dwell time relative boot-strap error, % |
| **MesoRD GT** | 45,3 | | 2,0 | | | 54,7 | | 2,4 | | |
| **tracked GT** | 41,1 | | 2,0 | | | 58,9 | | 2,8 | | |
| **0 nm** | 43,3 | 0,6 | 8,6 | 0,4 | 4,3 | 56,7 | 0,6 | 9,9 | 0,4 | 4,0 |
| **25 nm** | 51,3 | 0,5 | 5,7 | 0,2 | 3,3 | 48,7 | 0,5 | 6,6 | 0,2 | 3,5 |
| **50 nm** | 34,1 | 0,6 | 5,3 | 0,3 | 5,4 | 65,9 | 0,6 | 9,9 | 0,4 | 4,5 |
| **75 nm** | 30,2 | 0,7 | 9,2 | 1,1 | 12,2 | 69,8 | 0,7 | 12,5 | 0,9 | 7,2 |
| **100 nm** | 36,0 | 0,6 | 6,0 | 0,3 | 5,6 | 64,0 | 0,6 | 10,9 | 0,5 | 4,3 |

| M/C: gamma/exp dwell time 2 s | | | | | | | | | | |
|---|---|---|---|---|---|---|---|---|---|---|
| | M occ., % | M occ. boot-strap error | M dwell time, s | M dwell time boot-strap error | M dwell time relative boot-strap error, % | C occ., % | C occ. boot-strap error | C dwell time, s | C dwell time boot-strap error | C dwell time relative boot-strap error, % |
| **MesoRD GT** | 45,7 | | 2,0 | | | 54,3 | | 2,4 | | |
| **tracked GT** | 40,7 | | 2,0 | | | 59,3 | | 2,8 | | |
| **0 nm** | 45,0 | 0,6 | 6,8 | 0,3 | 3,7 | 55,0 | 0,6 | 8,8 | 0,3 | 3,9 |
| **25 nm** | 50,5 | 0,5 | 4,0 | 0,1 | 2,6 | 49,5 | 0,5 | 4,4 | 0,1 | 2,6 |
| **50 nm** | 42,3 | 0,6 | 3,9 | 0,1 | 2,8 | 57,7 | 0,6 | 5,2 | 0,2 | 3,0 |
| **75 nm** | 43,2 | 0,6 | 3,6 | 0,1 | 2,8 | 56,8 | 0,6 | 5,3 | 0,2 | 3,0 |
| **100 nm** | 43,1 | 0,6 | 4,4 | 0,1 | 3,2 | 56,9 | 0,6 | 5,9 | 0,2 | 3,3 |

| M/C: gamma/gamma dwell time 2 s | | | | | | | | | | |
|---|---|---|---|---|---|---|---|---|---|---|
| | M occ., % | M occ. boot-strap error | M dwell time, s | M dwell time boot-strap error | M dwell time relative boot-strap error, % | C occ., % | C occ. boot-strap error | C dwell time, s | C dwell time boot-strap error | C dwell time relative boot-strap error, % |
| **MesoRD GT** | 45,3 | | 2,0 | | | 54,7 | | 2,4 | | |
| **tracked GT** | 41,4 | | 1,9 | | | 58,6 | | 2,6 | | |
| **0 nm** | 43,8 | 0,5 | 5,7 | 0,2 | 3,5 | 56,2 | 0,5 | 7,2 | 0,3 | 3,6 |
| **25 nm** | 49,6 | 0,5 | 3,4 | 0,1 | 2,6 | 50,4 | 0,5 | 4,1 | 0,1 | 2,4 |
| **50 nm** | 46,6 | 1,3 | 2,5 | 0,2 | 9,0 | 53,4 | 1,3 | 4,0 | 0,2 | 4,3 |
| **75 nm** | 49,8 | 1,4 | 2,6 | 0,3 | 10,2 | 50,2 | 1,4 | 3,9 | 0,2 | 4,6 |
| **100 nm** | 44,1 | 0,6 | 4,0 | 0,1 | 2,9 | 55,9 | 0,6 | 5,3 | 0,2 | 3,1 |

| M/C: exp/exp dwell time 4 s | | | | | | | | | | |
|---|---|---|---|---|---|---|---|---|---|---|
| | M occ., % | M occ. boot-strap error | M dwell time, s | M dwell time boot-strap error | M dwell time relative boot-strap error, % | C occ., % | C occ. boot-strap error | C dwell time, s | C dwell time boot-strap error | C dwell time relative boot-strap error, % |
| **MesoRD GT** | 48,5 | | 4,1 | | | 51,5 | | 4,4 | | |
| **tracked GT** | 44,2 | | 3,9 | | | 55,8 | | 5,0 | | |
| **0 nm** | 44,3 | 0,4 | 11,9 | 0,4 | 3,3 | 55,7 | 0,4 | 15,5 | 0,6 | 3,7 |
| **25 nm** | 51,9 | 0,4 | 6,4 | 0,2 | 2,9 | 48,1 | 0,4 | 7,9 | 0,3 | 3,2 |
| **50 nm** | 34,3 | 0,5 | 10,7 | 0,5 | 4,7 | 65,7 | 0,5 | 20,1 | 0,8 | 3,9 |
| **75 nm** | 34,8 | 0,4 | 11,9 | 0,6 | 4,9 | 65,2 | 0,4 | 23,1 | 1,1 | 4,6 |
| **100 nm** | 35,9 | 0,4 | 12,9 | 0,6 | 4,8 | 64,1 | 0,4 | 25,9 | 1,2 | 4,4 |

| M/C: gamma/exp dwell time 4 s | | | | | | | | | | |
|---|---|---|---|---|---|---|---|---|---|---|
| | M occ., % | M occ. boot-strap error | M dwell time, s | M dwell time boot-strap error | M dwell time relative boot-strap error, % | C occ., % | C occ. boot-strap error | C dwell time, s | C dwell time boot-strap error | C dwell time relative boot-strap error, % |
| **MesoRD GT** | 47,7 | | 4,0 | | | 52,3 | | 4,4 | | |
| **tracked GT** | 42,4 | | 3,6 | | | 57,6 | | 5,0 | | |
| **0 nm** | 46,2 | 0,4 | 8,1 | 0,2 | 3,0 | 53,8 | 0,4 | 11,2 | 0,4 | 3,2 |
| **25 nm** | 52,9 | 0,4 | 8,3 | 0,2 | 2,9 | 47,1 | 0,4 | 7,9 | 0,2 | 2,9 |
| **50 nm** | 36,5 | 0,4 | 6,6 | 0,2 | 3,6 | 63,5 | 0,4 | 11,7 | 0,4 | 3,4 |
| **75 nm** | 47,6 | 0,4 | 6,5 | 0,2 | 2,4 | 52,4 | 0,4 | 7,8 | 0,2 | 2,5 |
| **100 nm** | 36,9 | 0,4 | 10,0 | 0,5 | 4,8 | 63,1 | 0,4 | 15,8 | 0,6 | 4,0 |

| M/C: gamma/gamma dwell time 4 s | | | | | | | | | | |
|---|---|---|---|---|---|---|---|---|---|---|
| | M occ., % | M occ. boot-strap error | M dwell time, s | M dwell time boot-strap error | M dwell time relative boot-strap error, % | C occ., % | C occ. boot-strap error | C dwell time, s | C dwell time boot-strap error | C dwell time relative boot-strap error, % |
| **MesoRD GT** | 47,2 | | 4,0 | | | 52,8 | | 4,4 | | |
| **tracked GT** | 38,2 | | 3,6 | | | 61,8 | | 5,2 | | |
| **0 nm** | 41,1 | 0,4 | 7,7 | 0,2 | 3,1 | 58,9 | 0,4 | 10,4 | 0,3 | 2,8 |
| **25 nm** | 49,6 | 0,4 | 8,0 | 0,2 | 3,1 | 50,4 | 0,4 | 7,5 | 0,2 | 2,6 |
| **50 nm** | 35,1 | 0,4 | 7,1 | 0,2 | 3,3 | 64,9 | 0,4 | 9,5 | 0,2 | 2,6 |
| **75 nm** | 36,8 | 0,4 | 7,6 | 0,3 | 3,3 | 63,2 | 0,4 | 9,3 | 0,3 | 2,9 |
| **100 nm** | 37,3 | 0,4 | 8,9 | 0,3 | 3,3 | 62,7 | 0,4 | 10,2 | 0,3 | 3,0 |

| M/C: exp/exp dwell time 8 s | | | | | | | | | | |
|---|---|---|---|---|---|---|---|---|---|---|
| | M occ., % | M occ. boot-strap error | M dwell time, s | M dwell time boot-strap error | M dwell time relative boot-strap error, % | C occ., % | C occ. boot-strap error | C dwell time, s | C dwell time boot-strap error | C dwell time relative boot-strap error, % |
| **MesoRD GT** | 49,5 | | 8,3 | | | 50,5 | | 8,7 | | |
| **tracked GT** | 49,7 | | 7,7 | | | 50,3 | | 9,6 | | |
| **0 nm** | 53,3 | 0,4 | 27,5 | 1,1 | 4,0 | 46,7 | 0,4 | 22,4 | 0,8 | 3,6 |
| **25 nm** | 46,5 | 0,3 | 13,1 | 0,4 | 3,0 | 53,5 | 0,3 | 16,3 | 0,5 | 3,1 |
| **50 nm** | 48,8 | 0,3 | 13,7 | 0,4 | 2,9 | 51,2 | 0,3 | 16,5 | 0,5 | 2,9 |
| **75 nm** | 46,8 | 0,4 | 20,2 | 0,7 | 3,4 | 53,2 | 0,4 | 26,5 | 1,0 | 3,6 |
| **100 nm** | 47,9 | 0,4 | 19,3 | 0,7 | 3,5 | 52,1 | 0,4 | 29,6 | 1,2 | 3,9 |

| M/C: gamma/exp dwell time 8 s | | | | | | | | | | |
|---|---|---|---|---|---|---|---|---|---|---|
| | M occ., % | M occ. boot-strap error | M dwell time, s | M dwell time boot-strap error | M dwell time relative boot-strap error, % | C occ., % | C occ. boot-strap error | C dwell time, s | C dwell time boot-strap error | C dwell time relative boot-strap error, % |
| **MesoRD GT** | 49,4 | | 8,0 | | | 50,6 | | 8,7 | | |
| **tracked GT** | 52,8 | | 6,2 | | | 47,2 | | 10,0 | | |
| **0 nm** | 55,0 | 0,3 | 12,4 | 0,3 | 2,4 | 45,0 | 0,3 | 24,3 | 0,9 | 3,7 |
| **25 nm** | 47,7 | 0,3 | 9,7 | 0,2 | 2,3 | 52,3 | 0,3 | 26,7 | 0,9 | 3,3 |
| **50 nm** | 50,0 | 0,3 | 10,6 | 0,2 | 2,3 | 50,0 | 0,3 | 25,5 | 0,9 | 3,4 |
| **75 nm** | 51,1 | 0,3 | 10,5 | 0,2 | 2,2 | 48,9 | 0,3 | 24,2 | 0,8 | 3,4 |
| **100 nm** | 45,4 | 0,3 | 14,4 | 0,5 | 3,2 | 54,6 | 0,3 | 40,2 | 2,4 | 5,9 |

| M/C: gamma/gamma dwell time 8 s | | | | | | | | | | |
|---|---|---|---|---|---|---|---|---|---|---|
| | M occ., % | M occ. boot-strap error | M dwell time, s | M dwell time boot-strap error | M dwell time relative boot-strap error, % | C occ., % | C occ. boot-strap error | C dwell time, s | C dwell time boot-strap error | C dwell time relative boot-strap error, % |
| **MesoRD GT** | 47,9 | | 7,5 | | | 52,1 | | 8,6 | | |
| **tracked GT** | 46,5 | | 5,8 | | | 53,5 | | 14,8 | | |
| **0 nm** | 49,5 | 0,3 | 9,7 | 0,2 | 2,3 | 50,5 | 0,3 | 25,8 | 0,9 | 3,6 |
| **25 nm** | 54,2 | 0,3 | 7,9 | 0,2 | 2,5 | 45,8 | 0,3 | 13,9 | 0,5 | 3,3 |
| **50 nm** | 41,4 | 0,3 | 10,6 | 0,3 | 2,5 | 58,6 | 0,3 | 40,1 | 1,5 | 3,8 |
| **75 nm** | 45,0 | 0,3 | 9,4 | 0,2 | 2,3 | 55,0 | 0,3 | 31,1 | 1,2 | 3,8 |
| **100 nm** | 40,8 | 0,3 | 12,1 | 0,4 | 3,1 | 59,2 | 0,3 | 49,5 | 2,6 | 5,2 |

### Supplementary Note 3. Convergence of HMM models

The size of the dataset determines how well the parameters can be estimated by HMM. To test how much data is required to reach the desired precision, the analysis can be performed on datasets of different sizes and the convergence of the model parameters can be followed. Previously we have investigated the convergence of a simulated model with cumulative datasets (see for example Fig. S18 in main text reference 3 and Fig. 5c in main text reference 17) which can be also performed on experimental data by selective inclusion of trajectories.

To produce the convergence plot, the order of the total pool of trajectories was first randomized. Then, for each HMM calculation, only the trajectories up to a specific index were included, such that every new subset is computed with the amount of data increasing in a cumulative manner by a factor 1.5. The same procedure was repeated 10 times with new randomizations of the total trajectory pool, giving rise to funnel-like convergence plots consisting of 10 curves for every parameter (CFE, occupancy, and dwell time) (Fig. S7).

We investigated the convergence of models with gamma distributed dwell times for both M and C states, and dwell times from 1 – 8 s (Fig. S7 and Table S2). We found that for models with dwell times 1 – 4 s, the variation in occupancy obtained with the maximum tested amount of data is within ±1 percent unit (approximately 1-3% relative standard deviation (RSD)) and dwell time is within ±0.2 s (less than 8% RSD) from the final mean value. For the 8 s model, convergence is slower. Here the occupancy has converged within ±6 percent units (approximately 12% RSD) with the complete dataset, whereas the dwell time for M state has converged within ±1 s (11% RSD). Dwell times for the C state converged much more poorly, within ±12 s (57% RSD).

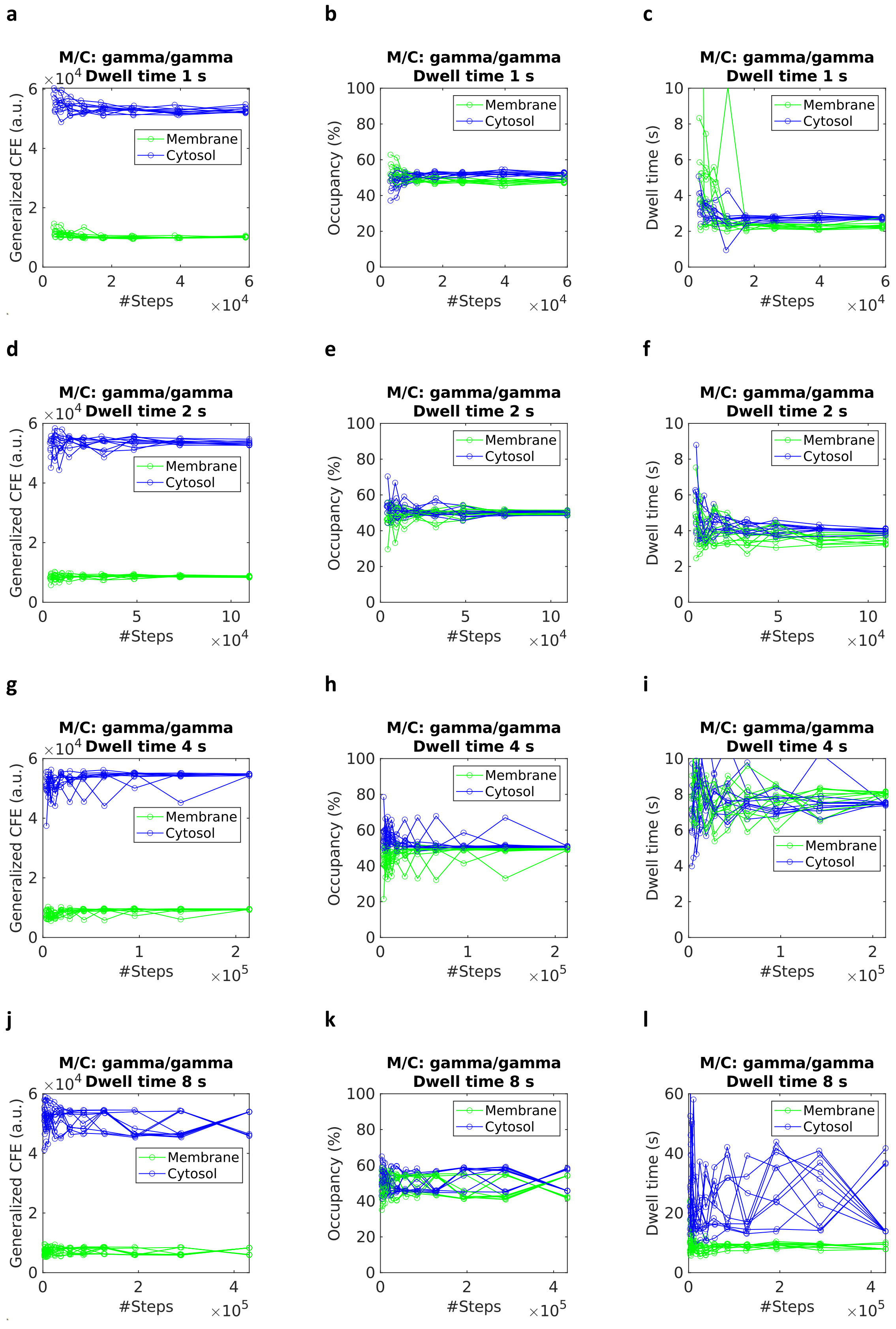

**Figure S7**. The convergence for 8-state HMM models coarse grained to 2 states. Generalized CFE (**a, d, g, j**), occupancy (**b, e, h, k**) and dwell time (**c, f, i, l**).

**Table S2.** Results of convergence for models with gamma distributed dwell times for M and C states. Standard deviation (STD) of mean final values and relative standard deviation (RSD) are also computed.

| | **M state, mean final value** | **M state, STD** | **M state, RSD, %** | **C state, mean final value** | **C state, STD** | **C state, RSD, %** |
|---|---|---|---|---|---|---|
| **Dwell time 1 s** | | | | | | |
| **Generalized CFE** | 10128 | 208 | 2.1 | 52908 | 941 | 1.8 |
| **Occupancy, %** | 48 | 1 | 2,7 | 52 | 1 | 2.6 |
| **Dwell time, s** | 2.3 | 0.2 | 7.8 | 2.7 | 0.1 | 2.2 |
| **Dwell time 2 s** | | | | | | |
| **Generalized CFE** | 8608 | 214 | 2.5 | 53353 | 674 | 1.3 |
| **Occupancy, %** | 50 | 1 | 2.1 | 50 | 1 | 2.0 |
| **Dwell time, s** | 3.5 | 0.2 | 7.0 | 4.0 | 0,1 | 3.2 |
| **Dwell time 4 s** | | | | | | |
| **Generalized CFE** | 9486 | 113 | 1,2 | 54656 | 267 | 0.5 |
| **Occupancy, %** | 49 | 0 | 0,7 | 51 | 0 | 0.6 |
| **Dwell time, s** | 8.0 | 0.2 | 2,5 | 7.5 | 0.1 | 0.9 |
| **Dwell time 8 s** | | | | | | |
| **Generalized CFE** | 7635 | 1128 | 14,8 | 51606 | 3775 | 7.3 |
| **Occupancy, %** | 51 | 6 | 11.7 | 49 | 6 | 12 |
| **Dwell time, s** | 8.4 | 0.9 | 10.6 | 21 | 12 | 56 |

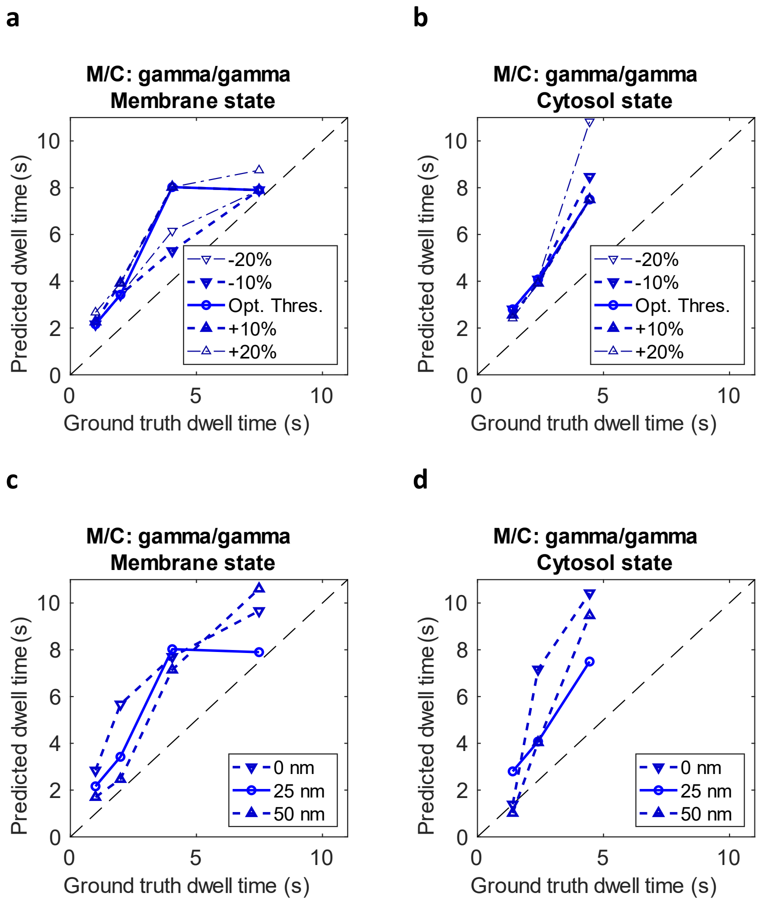

**Figure S8.** The predicted dwell time plotted against the ground truth dwell time for membrane state (**a, c**) and cytosol state (**b, d**) for models with dwell time of 1, 2, 4 and 8 s for membrane state and 1, 2 and 4 s for cytosol state. The dwell time of both the membrane and cytosol state is gamma distributed. The threshold used to coarse grain the 8-state models was varied by either ±10% or ±20% from the selected threshold (**a, b**) or the allowed circle fit moving distances used was 0, ±25 and ±50 nm (**c, d**).

**Table S3.** Dwell times obtained for models with gamma distributed M and C dwell times of 1 – 8 s with varying thresholds and circle moving distances. Standard deviations (STD) and relative standard deviation (RSD) are also computed.

| **M/C: gamma/gamma Membrane state** | | | | | |
|---|---|---|---|---|---|
| | 1 s | 2 s | 4 s | 8 s | |
| | 10.1 | 8.0 | 16.1 | 4.8 | **RSD, %** |
| | 0.2 | 0.3 | 1.3 | 0.4 | **STD** |
| **-20%** | 2.2 | 3.4 | 6.2 | 7.9 | **Detected dwell times, s** |
| **-10%** | 2.2 | 3.4 | 5.3 | 7.9 | |
| **selected threshold** | 2.2 | 3.4 | 8.0 | 7.9 | |
| **+10%** | 2.3 | 3.9 | 8.0 | 7.9 | |
| **+20%** | 2.7 | 3.9 | 8.0 | 8.7 | |

| **M/C: gamma/gamma Cytosol state** | | | | | |
|---|---|---|---|---|---|
| | 1 s | 2 s | 4 s | 8 s | |
| | 6.4 | 2.0 | 19.2 | | **RSD, %** |
| | 0.2 | 0.1 | 1.4 | | **STD** |
| **-20%** | 2.8 | 4.1 | 10.8 | | **Detected dwell times, s** |
| **-10%** | 2.8 | 4.1 | 8.5 | | |
| **selected threshold** | 2.8 | 4.1 | 7.5 | | |
| **+10%** | 2.6 | 3.9 | 7.5 | | |
| **+20%** | 2.4 | 3.9 | 7.5 | | |

| **M/C: gamma/gamma Membrane state** | | | | | |
|---|---|---|---|---|---|
| | 1 s | 2 s | 4 s | 8 s | |
| | 26.7 | 48.0 | 5.6 | 17.4 | **RSD, %** |
| | 0.6 | 1.6 | 0.4 | 1.4 | **STD** |
| **0 nm** | 2.8 | 5.7 | 7.7 | 9.7 | **Detected dwell times, s** |
| **25 nm** | 2.2 | 3.4 | 8.0 | 7.9 | |
| **50 nm** | 1.7 | 2.5 | 7.1 | 10.6 | |

| **M/C: gamma/gamma Cytosol state** | | | | | |
|---|---|---|---|---|---|
| | **1 s** | **2 s** | **4 s** | **8 s** | |
| | 33.6 | 44.0 | 19.9 | | **RSD, %** |
| | 0.9 | 1.8 | 1.5 | | **STD** |
| **0 nm** | 1.4 | 7.2 | 10.4 | | **Detected dwell times, s** |
| **25 nm** | 2.8 | 4.1 | 7.5 | | |
| **50 nm** | 1.0 | 4.0 | 9.5 | | |
